# Signalling-dependent refinement of cell fate choice during tissue remodelling

**DOI:** 10.1101/2023.02.21.529250

**Authors:** Sophie Herszterg, Marc de Gennes, Simone Cicolini, Anqi Huang, Cyrille Alexandre, Matthew Smith, Helena Araujo, Jean-Paul Vincent, Guillaume Salbreux

**Author notes:** Equivalent contribution (Sophie Herszterg, Marc de Gennes, Simone Cicolini). Equivalent senior authors for correspondence (Jean-Paul Vincent, Guillaume Salbreux). Address correspondence to: Jean-Paul Vincent or Guillaume Salbreux.

## Abstract

How biological form emerges from cell fate decisions and tissue remodelling is a fundamental question in development biology. However, an understanding of how these processes operate side-by-side to set precise and robust patterns is largely missing. Here, we investigate this interplay during the process of vein refinement in the *Drosophila* pupal wing. By following reporters of signalling activity dynamically, together with tissue flows, we show that longitudinal vein refinement arises from a combination of local tissue deformation and cell fate adjustments controlled by a signalling network involving Notch, Dpp, and EGFR. Perturbing large-scale convergence and extension tissue flows does not affect vein refinement, showing that pre-patterned vein domains are able to intrinsically refine to the correct width. A minimal biophysical description taking into account key signalling interactions recapitulates the intrinsic tissue ability to establish a thin, regular vein independently of large-scale tissue flows. Supporting this prediction, artificial proveins optogenetically generated orthogonal to the axis of wing elongation refine against large-scale flows. Overall, we find that signalling-mediated updating of cell fate is a key contributor to reproducible patterning.

## INTRODUCTION

The development of functional tissues requires that specialized cells be generated in the right numbers and at the right place. This is achieved largely by spatio-temporal regulation of the genes that control cell differentiation, numbers and morphology. The precision and reproducibility of developmental gene expression patterns is remarkable, especially in dynamic tissues that grow and change shape as cells acquire their identities. One classic strategy is for fate specification to proceed by progressive improvements in spatial precision. Thus, cells acquire rough positional cues from morphogen gradients, and, as the tissue expands and changes shape, subsequent cell interactions and gene regulatory networks refine positional information to reach cellular resolution (reviewed in [1–7]).

In a tissue that undergoes large deformation, one would expect that the need to update cell fates is particularly acute. This is the case in *Drosophila* pupal wings, which undergo convergence and extension while veins are being specified from a rough pre-pattern. There, as already seen in the wing disc, the provein fate can be recognised by the absence of DSRF, a transcription factor that marks intervein territories [8, 9]. At the end of wing morphogenesis, the vein pattern is highly stereotypical, with each vein consisting of a DSRF-negative stripe of a width regularly alternating between 2 and 3 cells [10, 11]. In contrast, proveins, which are also recognised by the absence of DSRF in imaginal discs are broad with jagged and ill-defined boundaries [8, 9]. Thus, proveins are a rough template that is subsequently refined during pupal stages to give rise to a highly reproducible final vein pattern [12].

Vein refinement requires cell interactions mediated by a network involving multiple signalling pathways (reviewed in [13–15]). Although different combinations of factors initiate the formation of individual proveins, they all converge on the activation of EGFR signalling, which represses DSRF [9, 16–18]. Absence of this transcriptional repressor in turn allows the expression of *dpp* [19], which encodes a TGFβ, and *rhomboid*, which encodes an enzymatic activator of Spitz, an EGFR ligand [16, 20, 21]. Activation of Dpp and EGFR signalling further represses DSRF expression [18], triggering a feedback loop that reinforces the production of Dpp and Spitz by proveins. In response to EGFR signalling, provein cells also express Delta, which activates the Notch receptor in flanking intervein cells, thus preventing expansion of the provein territory [22, 23]. Indeed, gain of function of Notch signalling during imaginal or pupal development led to loss of vein tissue, while inactivation of Notch signalling results in the expansion of vein domains [18, 23, 24]. Therefore, provein cells produce vein-promoting and vein-inhibiting signals, which compete to specify cell fate. Although genetic manipulations show that the vein fate (as defined by the absence of DSRF) is plastic, it is not clear whether the above signalling network can account entirely for vein refinement since morphogenetic processes and/or lineage restrictions could also contribute [25, 26]. Notably, the cell flows that accompany tissue-wide convergence and extension could help provein domains to thin out [25, 27]. Additionally, vein cells have been shown to express higher levels of the adhesion molecule E-cadherin [10], suggesting that a cell sorting mechanism could be at play.

Here, we aim at disentangling the relative roles of tissue remodelling, signalling-mediated cell fate decision in ensuring refinement of the vein prepattern. We take advantage of pupal wings which are amenable to live imaging by confocal microscopy, opening the way to a dynamic understanding of cell fate refinement. With this aim in mind, we have created several fluorescent reporters to allow live tracking of signalling activity and cell fate choice. The resulting data, combined with mathematical modelling and optogenetics-based gain of function experiments allow us to conclude that provein refinement begins with local convergence and extension flows, followed by signalling-driven cell fate adjustments, and ends with additional refinement likely provided by cortical tension asymmetries. We highlight the importance of the signalling network involving Dpp, EGFR and Notch in ensuring reproducible patterning in the face of cell rearrangements and tissue morphogenesis. Furthermore, our minimal model of the signalling network, which involves opposing effects of long-range Notch-Delta signalling and short-range Dpp/EGF signalling, is sufficient to account for vein refinement and smoothening.

## RESULTS

### Provein refinement dynamics with a live reporter of cell fate

Fixed preparations of pupal wings show that proveins become progressively thinner and more sharply delineated as development proceeds [13]. To gain insight in the dynamics of provein refinement, we devised a live reporter of *DSRF* expression. DNA encoding green fluorescent protein (GFP, including a termination codon) was knocked in the *DSRF* locus immediately downstream of the translation initiation codon to generate a transcriptional reporter (DSRF>GFP, see Figure S1 and Methods for additional details). Live imaging of DSRF>GFP (a marker of intervein fate) confirmed that during the period from 16 to 34h APF, proveins start out broad and irregular and become progressively narrower and less rough (Figure 1A-C and Movie S1). Furthermore, comparing provein outlines across different individuals showed a high degree of variability at 16h APF and near congruence at 34h APF (Figure 1D and 1E), indicating that refinement allows proveins to converge to a reproducible structure.

**Figure 1:**
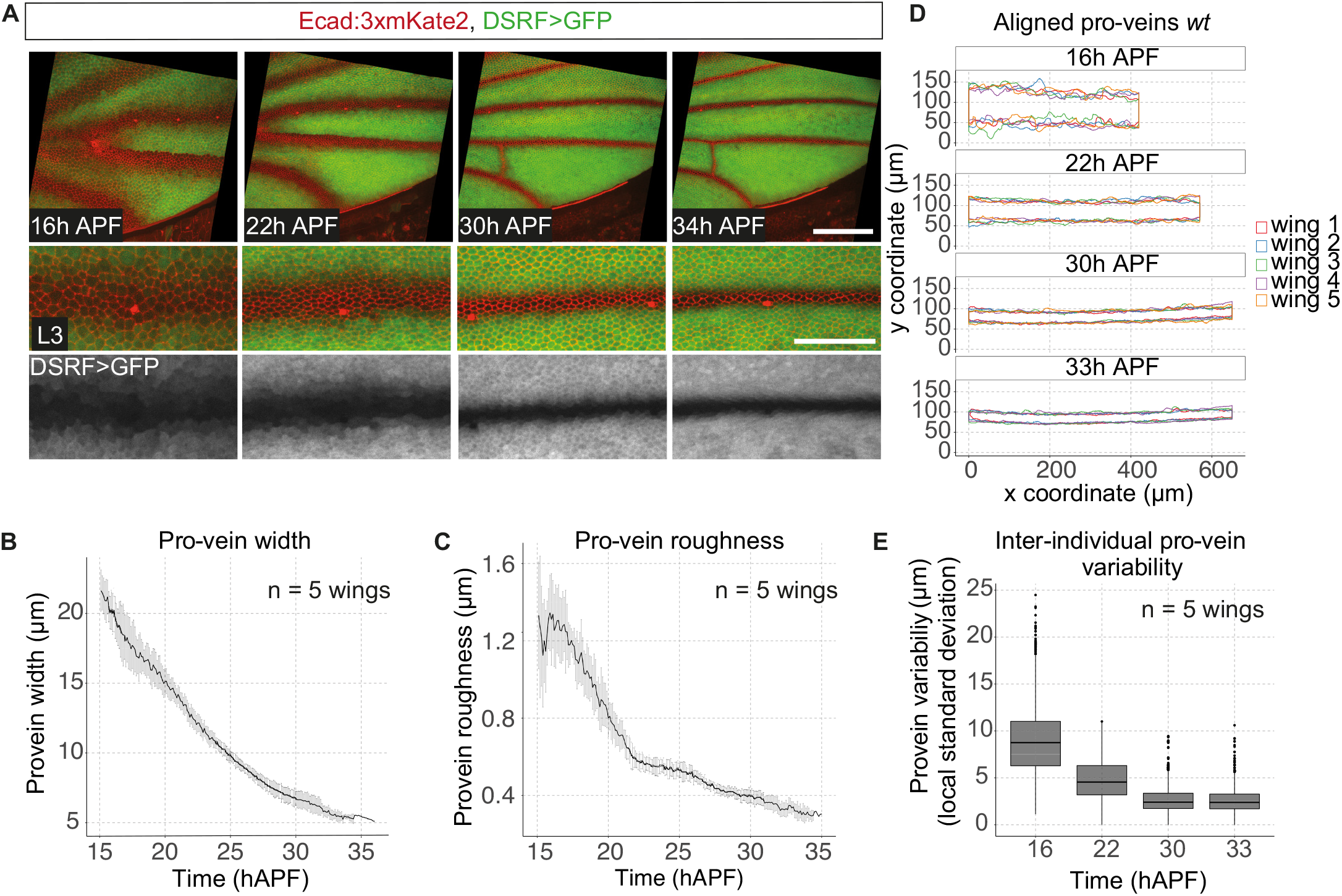
Dynamics of provein refinement in pupal wing. **A**. Top: timelapse images of a pupal wing expressing E-cadherin fused to 3xmKate2 (E-cad:3xmKate2, red) and the DSRF reporter (DSRF>GFP, green). Developmental time is indicated in hours after puparium formation (h APF). Bottom: close-ups of the L3 provein. **B**. Average L3 provein width over time. **C**. Average L3 provein roughness over time (see Methods for definition). **D**. Contours of the L3 provein from 5 wings, registered in space and time, from 16 to 33h APF. For C and D, n = 5 wings and shaded region indicates standard deviation. **E**. Boxplot showing L3 provein variability between 5 wings over time (see Methods for details). Horizontal lines correspond to the median, lower and upper box bounds to the first and third quartiles and whiskers to 1.5 times the inter-quartile range. Scale bars: 100 *μ*m (A, full wings) and 50 *μ*m (A, close-ups).

### Convergence and extension are not required for provein refinement

Provein refinement takes place between 16 and 34h APF (Figure 1A-C), a period when the pupal wing undergoes convergence and extension along the proximal-distal axis, parallel to longitudinal veins [27, 28]. To explore whether such tissue-scale rearrangements contribute to vein narrowing, we assessed DSRF>GFP dynamics in wings that are defective in convergence and extension. Dumpy (Dpy) is an apical extracellular matrix protein required for the pupal wing blade to anchor itself to the distal cuticle [27, 29, 30]. In the absence of Dpy and hence anchorage, convergence and extension are impaired, and the wing blade collapses toward the hinge [27, 29]. Nevertheless, as shown by tracking DSRF>GFP dynamics, *dpy* mutant proveins still narrowed (Figure 2A-C and Movie S2) and converged to a reproducible structure (Figure 2D and 2E). Mutant proveins also became as smooth as wild type veins, albeit with a small delay (Figure 2C). As expected from the near-normal behaviour of pupal proveins, adult veins appeared similar in thickness and shape in *dpy* mutant as in wild type wings (Figure S2). Since the dynamics of provein refinement in *dpy* is almost indistinguishable from that of wild type wings (Figure 2B), we conclude that tissue-scale convergence and extension of the pupal blade is not required for provein refinement.

**Figure 2:**
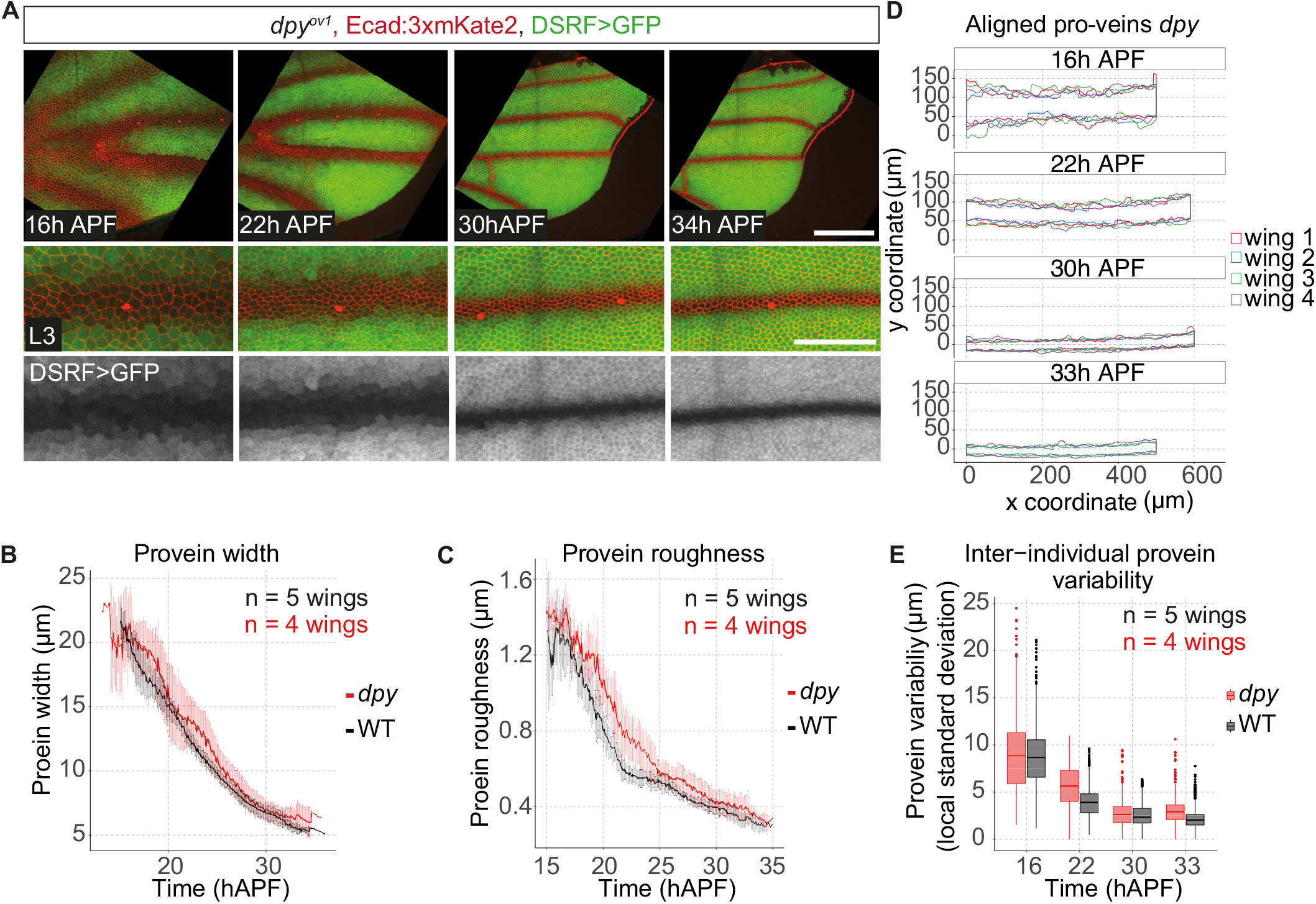
Dynamics of provein refinement in *dpy* pupal wing with perturbed convergence and extension. **A**. Top: timelapse images of a dumpy mutant (*dpy*) wing expressing E-cadherin fused to 3xmKate2 (E-cad:3xmKate2, red) and the DSRF reporter (DSRF>GFP, green). Developmental time is indicated in hours after puparium formation (h APF). Bottom: close-ups of the L3 vein. **B**. Average L3 provein width over time in wt (grey) and dpy mutant (red) wings. **C**. Average L3 provein roughness over time in wt (grey) and dpy mutant (red) wings. For B and C, n = 5 wings (wt), n = 4 wings (*dpy*) and shaded regions indicate standard deviation. **D**. Contours of the L3 proveins from 4 different dpy mutant wings, registered in space and time, from 16 to 33h APF. E. Boxplot of L3 provein variability between 5 wt wings (grey) and 4 dpy mutant wings (red) over time (see Methods for details). Horizontal lines correspond to the median, lower and upper box bounds to the first and third quartiles and whiskers to 1.5 times the inter-quartile range. In B, C and E the wt data shown is the same as in Figure 1, shown here to facilitate comparison. Scale bars: 100 *μ*m (A, full wings) and 50 *μ*m (A, close-ups).

### Cell fate adjustments play a major role in provein refinement

The results above show that provein refinement does not require tissue-wide cell flows and is therefore controlled more locally, within each provein. One possibility is that provein cells could undergo local shape changes or rearrangements that lead to narrowing of the whole domain. Alternatively, or in addition, refinement could follow from the loss of vein fate in a subset of provein cells (gain of DSRF expression). To assess the relative contribution of local cell shape change, cellular rearrangements and fate change to refinement, we used a cell contour marker (E-cadherin:mKatex3), along with our DSRF>GFP reporter, to track fate changes in all the cells classified as proveins at the onset of refinement (Figure 3A). Thus, we define the *initial* provein domain by the absence of GFP in the DSRF>GFP background at 16h APF, before the onset of refinement (see Methods and Figure S3A for details). In contrast, the *actual* provein is defined by the absence of GFP at the given time (Figure 3B). We then tracked the contour and GFP fluorescence of all the cells within the *initial* provein domain and their descendants until 32h APF, at the end of refinement (Figure 3A-D) (the initial provein’s progeny). This showed that only a subset the *initial* provein cells maintains the provein fate, both in wild type and *dpy* mutant wings (56%, 53% in wild type and 48%, 70% in *dpy*; note that the *dpy* mutant wing showing less refinement also had a thinner *initial* provein, Figure 3C). These ‘stable’ provein cells were located near the middle of the *initial* provein domain, while those that lost the provein fate tended to be located more peripherally (Figure 3D). This shows that modulation of cell fate adjustment is an important driver of provein refinement, in agreement with previous work with fixed preparations [13].

**Figure 3:**
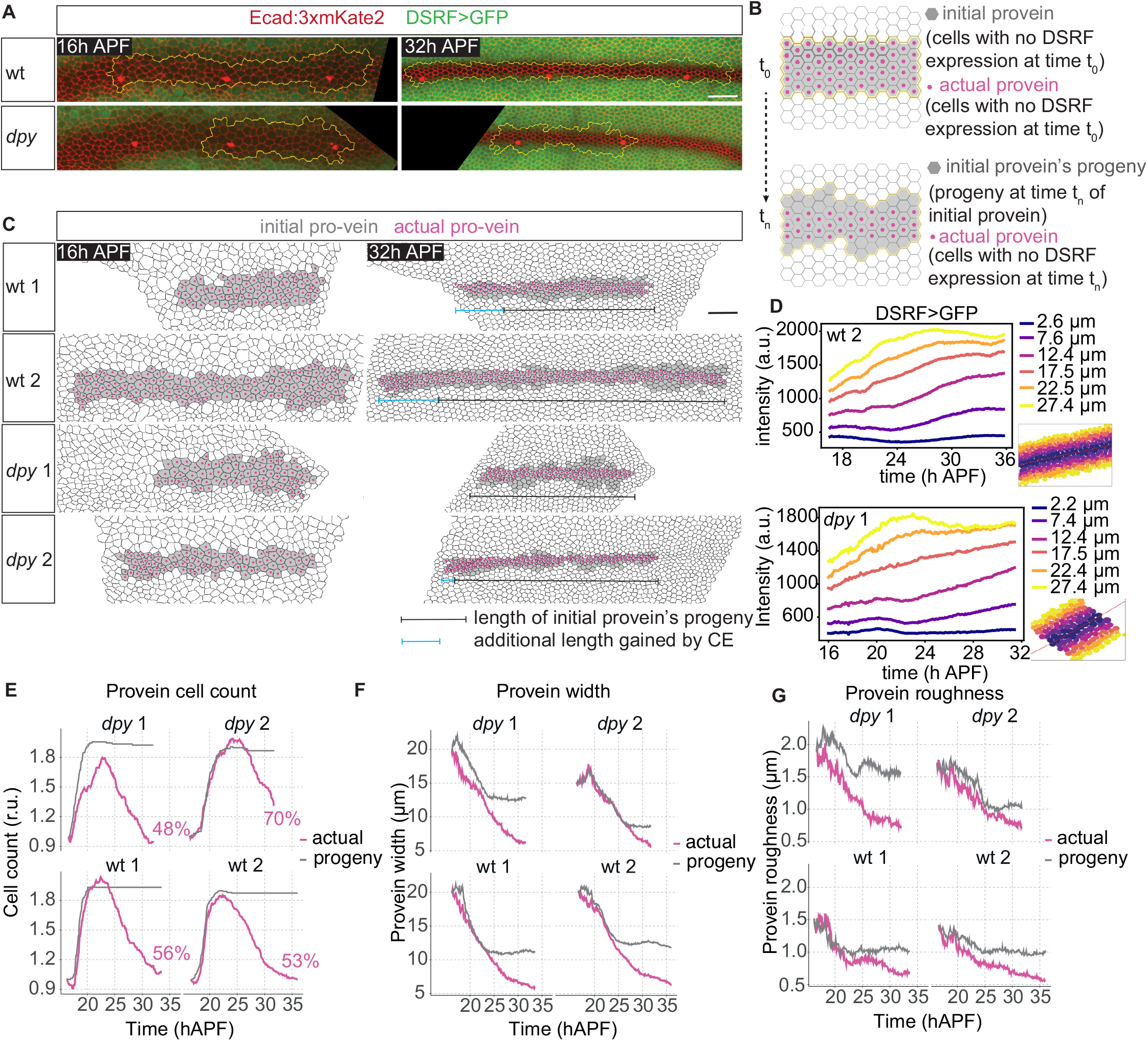
Cell fate adjustment plays a role in provein refinement. **A**. Timelapse images of the L3 vein of a wt (top) and *dpy* mutant (bottom) wing expressing E-cadherin fused to 3xmKate2 (E-cad:3xmKate2, red) and the DSRF reporter (DSRF>GFP, green). Cells within yellow contour: provein cells at 16h APF and their descendants at 32h APF. **B**. Schematic of the definition of the initial provein’s progeny domain (grey) and actual provein domain (magenta dots, see Methods for details) **C**. Timelapse images of the segmented L3 vein in 4 different animals (wt and *dpy* mutant) before and after refinement (16 and 32h APF respectively). Grey: initial provein’s progeny, magenta: actual provein (as defined in B). Black and blue bars: length of the initial provein’s progeny at 16h APF (black) and additional length gained by convergence and extension (CE, blue). Provein convergence and extension (CE) is severely affected in dpy mutant wings. **D**. Single-cell DSRF>GFP cell intensity over time and at different distances from the centre of the vein, in a wt (top) and a dpy mutant (bottom) wing. Distances to the centre of the provein are colour-coded from blue (closest) to yellow (farthest). Insets: positions of the cells that were quantified. See Figure S3F for examples in other wings. **E**. Number of cells belonging to the initial provein’s progeny (grey) and actual provein (magenta) over time in the L3 provein, for 4 individual wings. Values are normalised to the number of cells at the earliest developmental time. Percentage numbers: fraction of the initial provein’s progeny cells that contributed to the vein cells at the latest developmental time. **F**. L3 provein width over time, for 4 individual wings, for the initial provein’s progeny (grey) and actual provein (magenta) **G**. Roughness of the L3 provein over time (see Methods for details), for the initial provein’s progeny (grey) and actual provein (magenta). Scale bars: 20*μ*m (A and C).

To address when cell fate adjustment occurs, we compared the *actual provein* and *initial provein’s progeny* in terms of cell counts, provein width and roughness (Figure 3E-G). Between 16 and ~22h APF, as a result of a burst of wing-wide cell divisions [27], cell numbers increased in both the *actual provein* and *initial provein’s progeny* domains (Figure 3E). Beyond 22h APF, this proliferative activity ceased, leaving the number of cells within the *initial provein progeny* unchanged, while the number of *actual provein* cells (GFP-negative in DSRF>GFP) decreased, indicating that cell fate adjustments occur after 22h APF (Figure 3E). Provein width and roughness were also seen to decrease in two phases. During the first phase, lasting up to ~22h APF, the width and roughness of both initial provein’s progeny and actual proveins decreased concomitantly, suggesting an absence of cell fate change (cell fate change would be expected to alter the boundary of actual proveins relative to that of the initial provein’s progeny). We therefore attribute this phase of refinement to local tissue deformation mediated by cell shape changes and cellular rearrangements. In the second phase, the width and roughness of the initial provein’s progeny remained roughly constant while the actual provein domain became thinner and smoother. Therefore, this phase of refinement can be attributed to cell fate changes at the edge of the provein domain (Figure 3F and 3G). It appears therefore that vein refinement occurs first through tissue deformation, and then through cell fate change.

Smooth compartment boundaries (low roughness) have been attributed to increased junctional tension at these cellular interfaces [31]. We wondered if the decrease in provein roughness was similarly associated to increased junctional tension at the boundary between the intervein and the provein domains. Imaging Myosin II distribution with an anti-phospho-Myosin light chain antibody (anti-pMLC) did not reveal a clear enrichment at the provein boundaries at 24h APF (Figure S3B). We then measured the recoil velocity after laser-mediated junction ablation and compared the results for junctions within proveins, interveins, or at the interface between these two (referred below as border junctions) (Figure S3C). Provein and intervein cells were distinguished using a NRE>mcherry transgene, which is expressed in the intervein cells that flank proveins (Figure S3D and see next section). Up to 28h APF, junctional tension was similar at border junctions and intervein-intervein junctions. Junctional tension was slightly larger at border junctions than at vein-vein junctions at 16-22 h APF, although this difference disappeared between 24-28 h APF (Figure S3C). We further observed that, at 24 h APF, Myosin II was slightly enriched within the longitudinal proveins, but not at their border (Figure S3B). After refinement was complete, between 35 h APF and 40 h APF, recoil velocity became markedly larger (2.1-2.3 fold larger) at border junctions when compared to vein or intervein junctions (Figure S3C). Accordingly, the vein-intervein interface looked smoother at 35h APF (Figure S3E). These observations suggest that mechanical tension could contribute to the smoothening of the vein-intervein interface, albeit only at late stages, after the vein-intervein fates have been resolved.

As expected from our earlier observation that provein narrowing occurs near normally in *dpy* mutant wings, we found that provein cell count, width and roughness relaxed to similar values in *dpy* mutant wings as in wild type wings, despite the impaired tissue-wide convergence and extension flows (Figure 3E-G). The first phase of narrowing occurred at a similar rate and extent in *dpy* and wild type wings, indicating that the initial vein narrowing phase, attributed to local tissue deformation of the provein domain, is independent of convergence and extension, which occurs at the scale of the wing blade(Figure 3F). Although subsequent narrowing was more variable in *dpy* than in wild type, the provein domain reached the same width in the two genotypes at 32h APF. This suggests that cell fate changes can correct early defects to achieve a reproducible final width, highlighting the importance of cell fate adjustment for robustness.

### Signalling dynamics during provein refinement

To determine how cell fate changes adjust to the demands of tissue dynamics, we built on previous work, which showed that provein cells produce both vein-promoting and vein-inhibiting signals [13]. Provein cells express Dpp [19, 32], as well as *rhomboid*, which triggers the release of the EGFR ligand Spitz [16, 20, 33]. Thus, provein cells produce two diffusible signals that inhibit DSRF expression and could hence stimulate vein formation in surrounding cells. In addition, provein cells express Delta, a ligand of Notch, which, when activated, counteracts the inhibitory effect of EGFR signalling on *DSRF* expression. Accordingly, Notch signalling favours DSRF expression, enabling cells to acquire the intervein fate [21–23]. To understand how these three signalling pathways are integrated dynamically, we tracked signalling activity with live reporters during provein refinement.

We measured Notch signalling dynamics with a NRE>mcherry transgene [34] in the presence of our DSRF>GFP reporter to visualise proveins. Live imaging confirmed that Notch signalling is activated at both sides of the proveins (Figure 4A). At 16h APF, NRE>mcherry can be detected up to 8 cell diameters away from the proveins’ edges (Figure 4A) before becoming more restricted (Figure 4A–4B, and Movie S3). It appears therefore that during the early phase of refinement, Delta, which is only produced by provein cells, acts over several cell diameters (Figure S4A-C). Immuno-fluorescence revealed the presence of Delta punctae several cell diameters away from the proveins (Figure S4A), suggesting that Delta can physically spread, perhaps as a result of transport along basal protrusions, as reported in the fly notum (Figure S4B, S4C and [35, 36]). Plotting the activities of the DSRF and NRE>mcherry over time, and as a function of the distance from the vein center, showed that, as veins refine, an initially broad zone of Notch signalling activity on each side becomes narrower and progresses towards the center of the vein (Figure 4C and 4D). In addition, examination of single cell behavior indicated that activation of Notch precedes the downregulation of DSRF (Figure S4D). These data suggest that narrowing of Notch signalling at both sides of the proveins allow the intervein domain to encroach into proveins.

**Figure 4:**
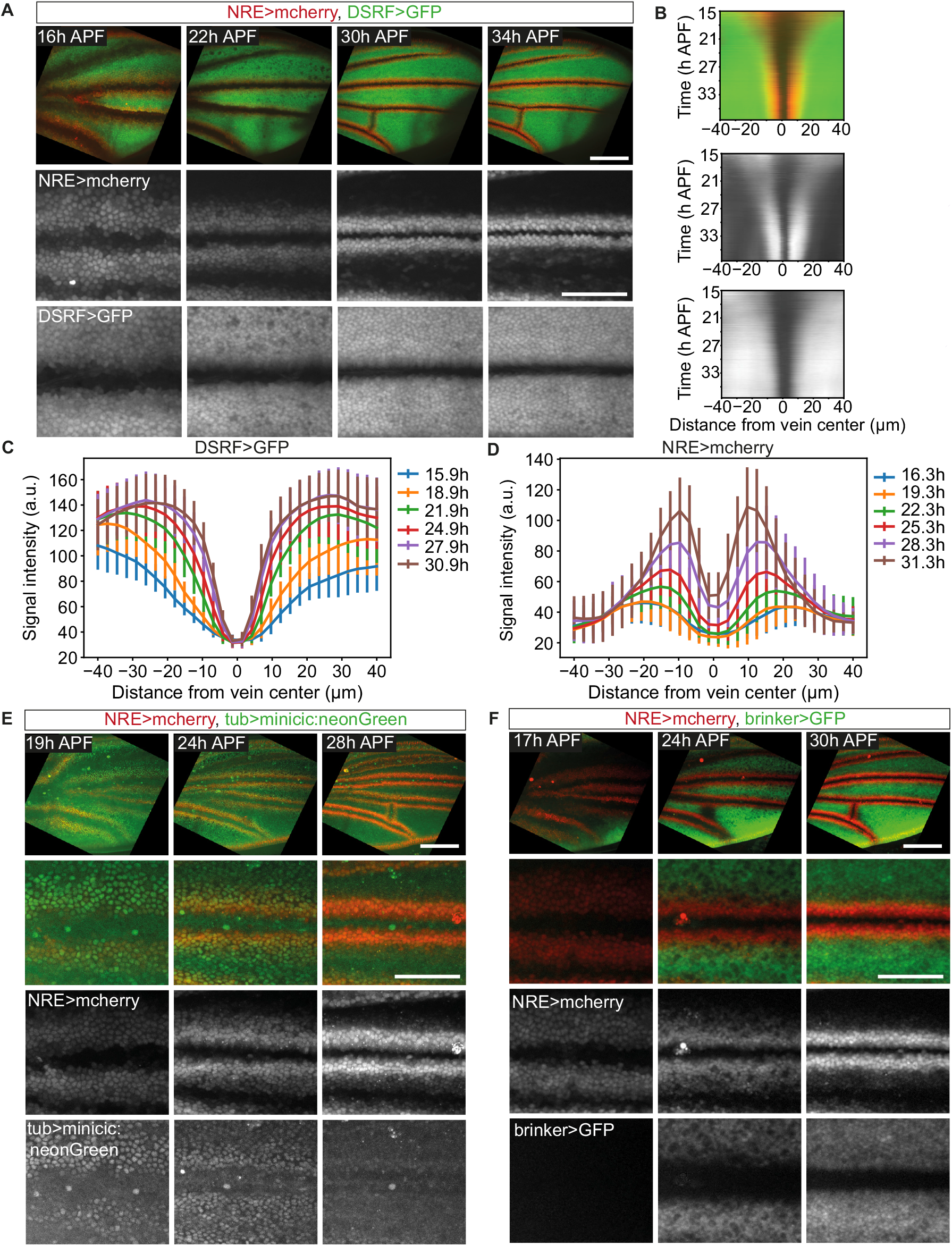
Signalling dynamics during provein refinement. **A**. Top: timelapse images of a pupal wing expressing a Notch signalling reporter (NRE>mcherry, red) and the DSRF expression reporter (DSRF>GFP, green). Bottom: close-ups of the L3 vein. **B**. Kymograph of fluorescence intensity across vein L3 (same wing as panel A). Notch signalling activity is depicted in red (NRE>mcherry) and DSRF expression in green (DSRF>GFP). Anterior is on the left. **C, D**. Average profiles of DSRF>GFP signal intensity (C) and Notch signalling (NRE>mcherry, D) around the center of the vein, at different times (n = 5 wings expressing DSRF>GFP and E-cad:3xmKate2 and n = 4 wings expressing NRE>mcherry and E-cad:GFP). Error bars indicate standard deviation. **E**. Top: timelapse images of a pupal wing expressing a Notch signalling reporter (NRE>mcherry, red) and a EGFR signalling reporter (tub>minicic:neonGreen, green). Absence of GFP signal indicates active EGFR signalling. Bottom: close-ups of the L3 vein. **F**. Top: timelapse images of a pupal wing expressing a Notch signalling reporter (NRE>mcherry, red) and a Dpp signalling reporter (brinker>GFP, green). Absence of GFP signal indicates active Dpp signalling. Close-ups show the L3 vein. Dpp signalling range decreases dramatically from 17 to 30h APF. Scale bars: 100 *μ*m (A, E and F, full wings) and 50 *μ*m (A, E and F, close-ups).

We next tracked the signalling pathways that <u>promote</u> the vein fate (Figure 4E and 4F). To assess EGFR signalling dynamics, we used a uniformly expressed Capicua-neonGreen fusion protein, which localises to the nucleus unless EGFR signalling is active (minicic-neonGreen, mScarlet version described in Ref. [37]). For Dpp signalling, we used a brinker>GFP (brk>GFP) transgene, which is repressed by signalling [38]. Throughout refinement, EGFR signalling was seen to be active in the provein as well as in adjoining intervein cells expressing NRE>mcherry (Figure 4E and Movie S4). Dpp signalling was initially (at 16h APF) active in the whole wing, presumably in response to long range action of Dpp previously produced at the A/P boundary. However after 24h APF, after Dpp has become expressed in proveins [32], signalling became restricted to within a few cell diameters from the edge of each provein (Figure 4F and Movie S5). A similar pattern of Dpp signalling activity was described previously from immunofluorescence analysis with an antibody against phosphorylated Mad [18, 39]. We conclude that, during refinement, the range of the two vein-promoting signals becomes relatively short.

Overall, our live imaging data confirm and extend previous analyses of signalling activity showing that provein cells produce both short-range vein-promoting signals and a long-range vein-inhibiting signal. To understand how this signalling network ensures robust and reproducible provein refinement, we formalised our observations in a mathematical model.

### Minimal model of signalling interactions leading to vein refinement

We developed a minimal description of the signalling network that controls cell fate adjustment in the pupal wing. Denoting *u_i_* a cell state variable that represents how close a cell *i* is to be in the vein (*u_i_* ≃ 1) or intervein (*u_i_* ≃ 0) state, we write (Figure 5A):

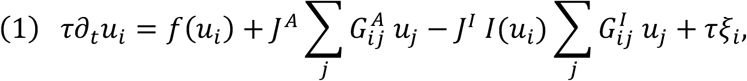

where in the right-hand side, the first term *f*(*u*) represents cell autonomous dynamics, the second term represents nearest neighbour activation with strength *J^A^*, the third term represents long range inhibition with strength *J^I^*, and the last term is a Langevin noise term with zero mean (Supplementary Theory). The autonomous dynamics is such that the cell state evolves according to a bistable potential *F*, defined by *f*(*u*) = −*∂_u_F*(*u*). Such bistable dynamics has been proposed to emerge from autocrine signalling mediated by Dpp and EGFR [40] and allows for a simple representation of cell fate decision [41]. Here we choose the potential *F* such that a parameter *r* controls the relative stability of the vein and intervein states (Supplementary Theory). We envision that short range activation arises from the combined action of Dpp and EGFR signalling, which we take to be short range (limited to nearest-neighbour interaction). This is encoded in the kernel 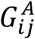. We also build on the observation that long-range inhibition by Notch-Delta signalling could be mediated by basal protrusions (Figure S4). We assume that the spatial range of Notch signalling activity by Delta, which is encoded in the kernel 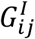, corresponds to an exponential decay on a distance of a few cells, in accordance with measurements of basal protrusion length at 20h APF (Figure S4C). The factor *I*(*u*) corresponds to inhibition of Notch signalling at high Delta level, as a result of cis-inhibition [42], ignoring for simplicity the non-uniform pattern of Notch expression [22, 23]. Eq. 1 constitutes a minimal model for the complex network of signalling interactions occurring in the process of vein refinement.

In the continuum limit where the distance between cells is small compared to other length scales in the system, and ignoring here Delta/Notch cis inhibition (*I* = 1), the equation for the cell state can be rewritten, in the absence of noise (Supplementary Theory):

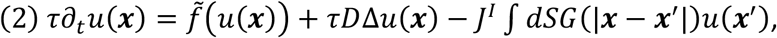

with *G* an exponentially decaying function of its argument, and the function 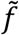 and parameter *D* are related to *f* and *J^A^*. The diffusion constant *D* is associated to nearest-neighbour vein state activation (arising from the term in*J^A^* in Eq. 1) and is an emerging quantity arising from cell-cell communication. Here, we choose 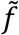 such that *u* = 0 and *u* = 1 are stable solutions in the absence of Notch inhibition, *J^I^* = 0. We use this continuum limit to obtain a phase diagram of steady-state solutions in two dimensions in the limit *J^A^* = 0 (Figure 5B). For *r* > 1/2, corresponding to the intervein state more stable, all cells are in the intervein state (Figure 5B and 5C). When the vein state is more stable (*r* < 1/2), at low values of the inhibition *J^I^*, the vein state invades the entire wing (Figure 5B), while for larger values of the inhibition parameter *J^I^*, localised stripes are solutions, indicating that the model can recapitulate vein-like patterns (Figure 5B and Figure S5, Supplementary Theory, [43]).

**Figure 5:**
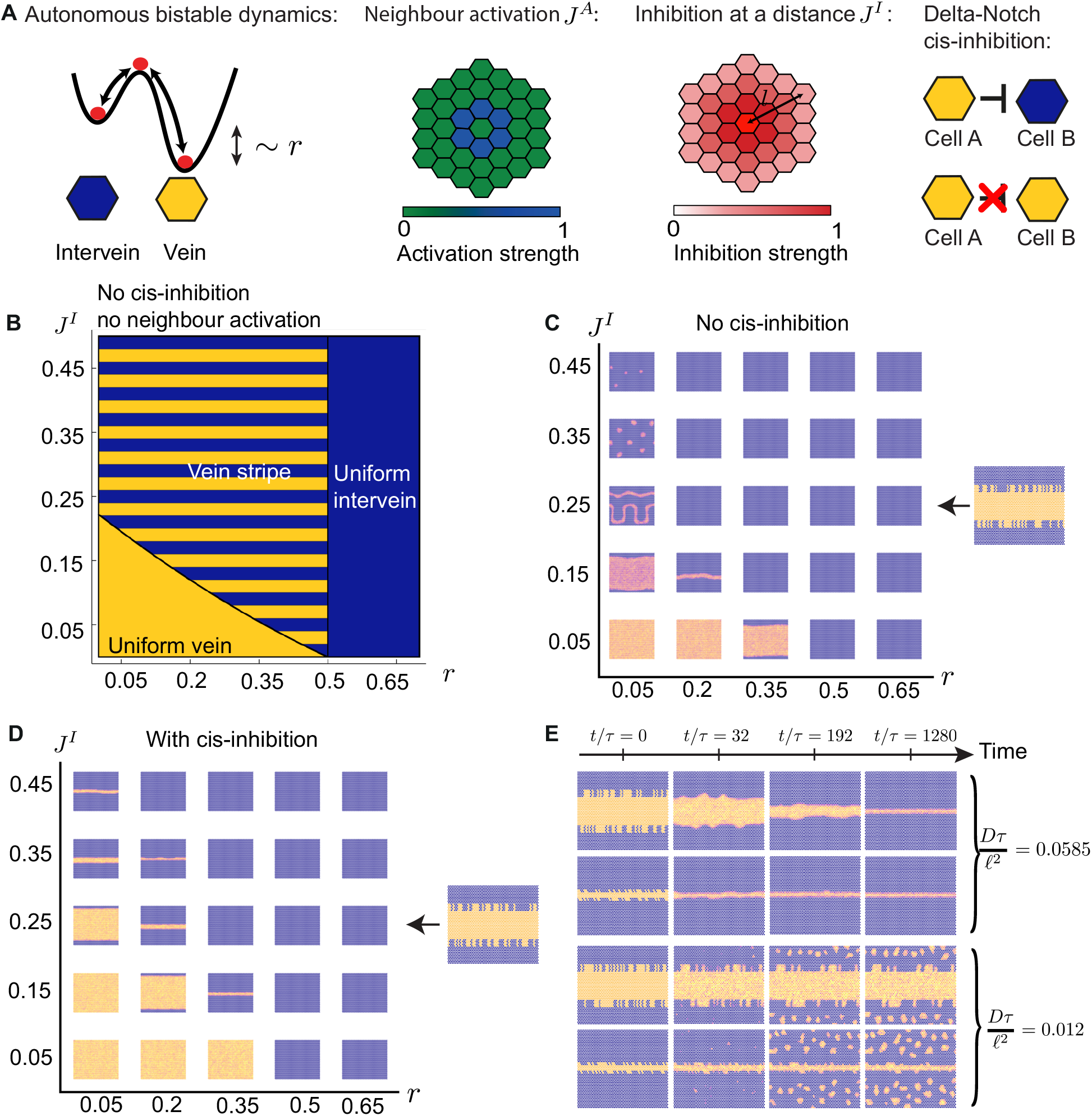
A minimal model of signalling interactions leading to vein refinement. **A**. Schematic of minimal model of signalling interactions. **B**. Approximate phase diagram indicating steady-state vein stripe solutions in the continuum limit and in the absence of near-neighbour activation (*J^A^* → 0, *D* → 0). **C**. Steady-state reached by model simulations on a hexagonal lattice (snapshots taken at *t/τ* = 1.4 × 10^4^), without Delta-Notch signalling cis-inhibition. The initial condition at *t* = 0 (right snapshot) corresponds to a broad noisy stripe. Other parameters: *τD/l^2^* = *3J^A^*Δ*x*^2^/(2*l*^2^) = 0.026, *γτ* = 0.0039, *l*/Δ*x* = 3 with Δ*x* the distance between nearest neighbours on the lattice. **D**. Steady-state reached by model simulations on an hexagonal lattice (snapshots taken at *t/τ* = 1.4 × 10^4^), with Delta-Notch signalling cis-inhibition. The initial condition at *t* = 0 (right snapshot) corresponds to a broad noisy stripe. The domain of existence of vein stripe solutions in parameter space is enlarged compared to panel C. *u_I_* = 0.52, *α_I_* = 0.075, other parameters as in C. See Supplementary Theory for details of parameter definitions. **E**. Simulation snapshots of relaxation of roughly defined provein domain towards a smoother vein domain, for two values of *τD/l^2^* = 3 *J^A^*Δ*x*^2^/(2*l*^2^). For high enough values of the diffusion constant D, the initial roughness in the vein pattern is suppressed. Other parameters: *r* = 0.1, *J^I^* = 0.4, *l*/Δ*x* = 2, *u_I_* = 0.52, *α_I_* = 0.075, *γτ* = 0.0039.

We then performed simulations of the original Eq. (1) without Delta-Notch cis-inhibition (*I* = 1) on a hexagonal lattice using a rough, thick stripe as initial condition, varying model parameters (Figure 5C and S5D-E). Nearest-neighbour activation, or equivalently a sufficiently large value of the diffusion constant *D*, is necessary to avoid ectopic activation of small vein domains triggered by noise and to smoothen irregular edges of the provein domain (Figure S5E). Indeed, at the level of the vein pattern, nearest-neighbour activation creates an effective line tension (not based on mechanics) acting on the boundary of a vein domain to straighten the interface (Supplementary Theory). This smoothening role of the effective diffusion arising from nearest-neighbour interactions also tends to eliminate the thin veins that appear at large values of *J^I^*, significantly reducing the parameter space in which stable veins appear (Figure 5C). In subsequent analysis, we found however that the vein-forming region is significantly enlarged when Delta-Notch cis-inhibition is taken into account (Figure 5D and S5F). Overall, incorporating essential features of known signalling interactions in a cell state model gives rise to a system that converges to a precise width and suppresses initial roughness at intermediate value of inhibition and for sufficiently high near-neighbour interactions (Figure 5E and Movie S6). This converging process is determined entirely from the dynamics of cell fate change arising from signalling interactions.

### Model of signalling interactions recapitulates on-the-fly provein refinement

We then proceeded to test if the model represented by Eq. 1 can predict refinement in the context of the complex morphogenetic flows that occur in the pupal wing. To mimic these flows, we used segmented movies of the pupal wing as templates for numerical simulations, where each tracked cell is allocated a value *u_i_* that changes with time (Figure 6A), and the nearest neighbour relations and distance between cells are updated at each time frame (Figure 6B). We also assume that, at cell division, the daughter cells inherit the state variable *u* of their mother. To compare the model with experimental measurements, we obtained an experimental value of *u* from the DSRF>GFP reporter fluorescence intensity through a linear mapping, assigning at each time point *u* = 0 to the cells with lowest DSRF>GFP fluorescence and *u* = 1 to the cells with highest DSRF>GFP fluorescence (Figure S6A, A’, Supplementary Theory). Initial conditions were set from experimental measurement at 16h APF, and boundary conditions were obtained by defining an outer boundary of segmented cells that maintain experimentally measured *u_i_* (Supplementary Theory). Importantly, such simulations test our model without making assumptions about cell flows and cell mechanics since cell motion is occurring exactly as in the biological system.

**Figure 6:**
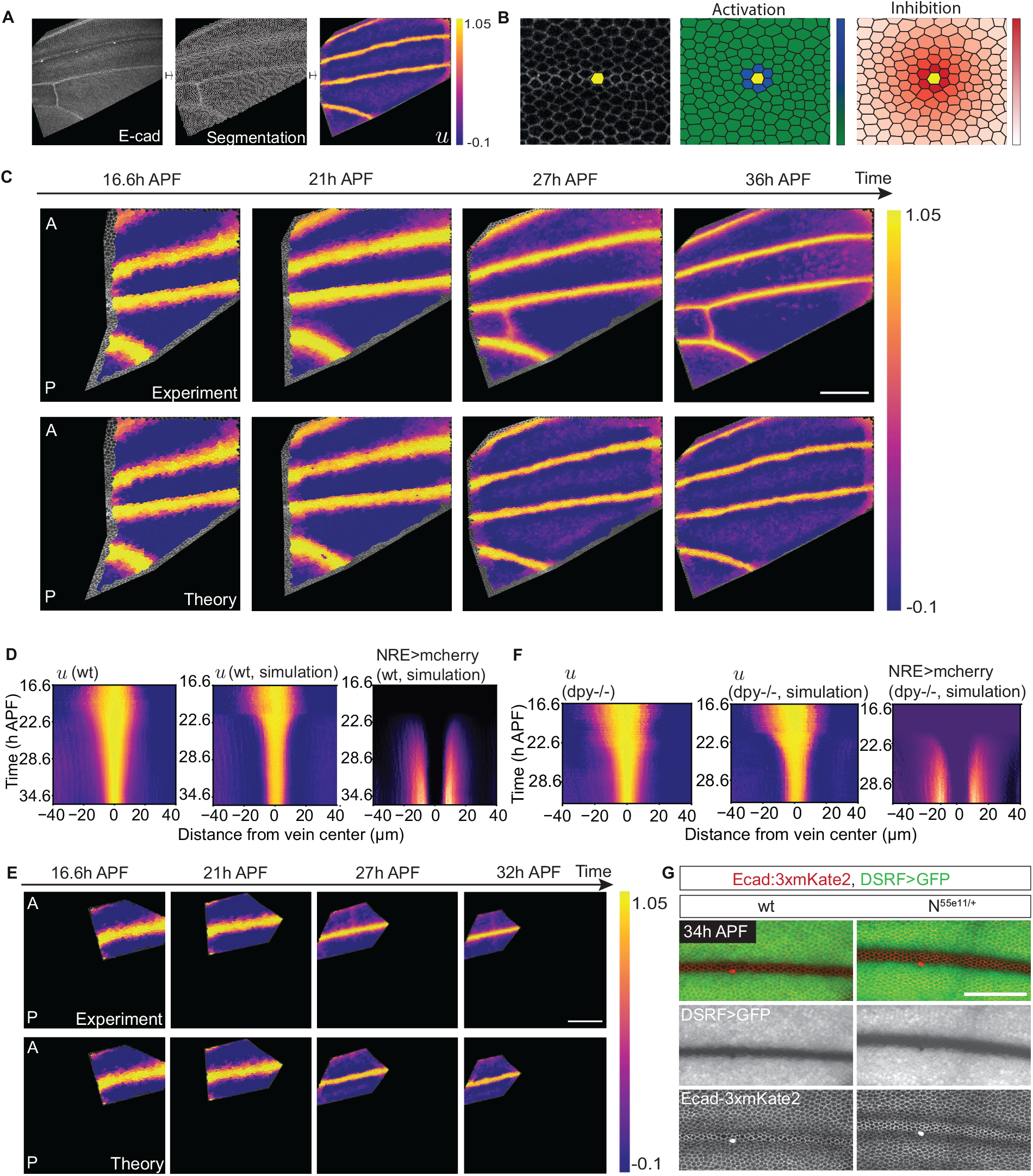
A minimal model of signalling interactions recapitulates vein refinement on a segmented wing template. **A**. Experimental measurement of DRSF intensity (left) is converted to a cell state variable *u* (right), with *u* = 0 corresponding to the intervein state and *u* = 1 to the vein state. **B**. Interaction rules in simulations on the segmented wing template. Each segmented cell (yellow, left) is allocated a value of the variable u, activates its nearest neighbours (middle) and inhibits its neighbours over a distance of a few cells (right). **C**. Snapshots showing the cell state variable derived from DSRF expression in a wt wing (top row, same wing as wt 2 in Figure 3), and cell state value u in a simulation on the wt segmented wing template (bottom row). Scale bar: 80μm. **D**. Kymographs of experimental cell state variable obtained from DRSF intensity, simulated cell state variable u, and simulated Notch activity. Same data as in panel C. Anterior is on the left. **E**. Snapshots showing the cell state variable derived from DSRF expression in a dpy mutant (top row, same wing as dpy 1 in Figure 3), and cell state dynamics obtained from a simulation on the dpy segmented wing template (bottom row). Scale bar: 80μm. **F**. Kymographs of experimental cell state variable obtained from DRSF intensity, simulated cell state variable *u*, and simulated Notch activity. Same data as in panel E. Anterior is on the left. **G**. Images of the refined L3 veins of wt (left) and Notch heterozygous mutant (N55e11/+, right) wings expressing E-cad:3xmKate2 and DSRF>GFP. Developmental time is indicated as hours after puparium formation (h APF). Scale bar: 50 μm. A: anterior, P: posterior. See Supplementary Theory for simulation parameters.

Parameters were then adjusted, using results on a hexagonal lattice as a guide (Figures 5 and S5F), to test whether vein refinement could be recapitulated. We assumed that between 16h APF and 21h APF cells keep their state variable values (Figure 6C), and that after 21h APF, cell states evolve dynamically according to Eq. 1 (Supplementary Theory). With these assumptions, we find that vein refinement can be captured to a large extent by our minimal model (Figure 6C, S6B,B’ and Movie S7). In both experiment and simulation, proveins were flanked by regions of Notch signalling activity (Figure 4B, 6D and S6). Experimental NRE>mcherry dynamics after 21h APF was recapitulated by the model by assuming a 20h decay time [44] for the reporter (Figure 4B, 6D and S6). Posterior cross-veins did not form in simulation, as expected since they are not present in the prepattern at 16h APF (Figure 6C). Consistent with our conclusion that refinement occurs through both tissue deformation and cell fate change, we found that merely advecting the cell fate variable (*du*/*dt* = 0) (i.e. cells move carrying a constant u) yielded incomplete vein refinement (Figure S6F,F’). Simulations were also implemented on a *dpy* mutant wing template and these similarly recapitulated experimental observations (Figure 6E and 6F), indicating that the simulated refinement process is insensitive to tissue-scale convergence extension along the proximal-distal axis. Although our simulations achieved significant refinement, real veins appeared slightly thinner and straighter than simulated ones (Figure 6C), suggesting that additional mechanisms are at play *in vivo*. As a further test of our model, we imaged heterozygous Notch mutant pupal wings, which are known to have reduced Notch signalling and thicker veins [24, 45]. As predicted by the model (Figure S6G,G’), L3 refinement was less complete than in wild-type wings (Figure 6G, Figure S6H-H”). Overall, we conclude that our minimal model of signalling interactions captures the essential features of provein refinement by cell fate change.

### Ectopic transverse veins refine but are fragile

Our model suggests that fate adjustments are sufficient to mediate provein refinement, even in the absence of mechanical contraction. It also predicts that signalling interactions alone can generate smooth stripes that converge to a reproducible width. We then wondered if, in our model, ectopic proveins could also refine. We introduced, at the start of the simulation, an ectopic transverse provein between two parallel proveins on a static hexagonal lattice (Figure S7A-B). Such transverse proveins narrowed down, either to reach a fixed width, or to disappear, depending on the parameters (Figure S7A-B). With parameters used to recapitulate provein refinement on the segmented wing template (Figure 6A-C), some virtual ectopic transverse proveins disappeared rapidly, while others remained up until 36h APF but disappeared over longer times, depending on the initial shape of the ectopic provein domain (Figure 7A). Therefore, in the model, transverse proveins refine but are more fragile and parameter-sensitive than longitudinal veins.

**Figure 7:**
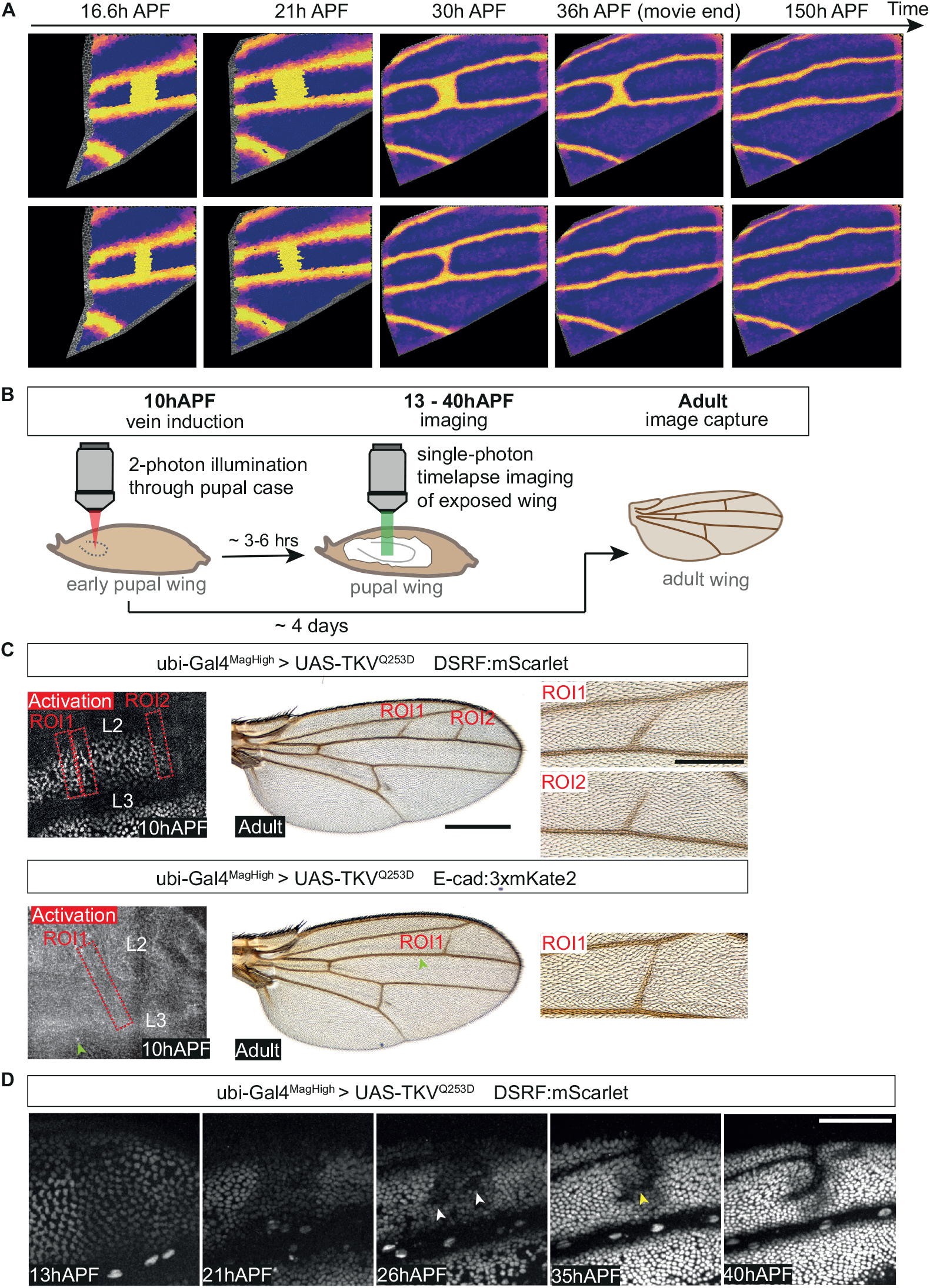
Ectopic transverse veins refine but are fragile. **A**. Snapshots of cell state dynamics obtained from simulation on a wt segmented wing template, introducing an additional tranverse vein domain in the initial condition between L3 and L4. Top and bottom rows: initial domains of different width. See Supplementary Theory for simulation parameters. **B**. Experimental setup used for the induction of ectopic veins by optogenetics (see Methods). Vein induction is performed by activating Shine-Gal4 in early pupal wings (10h APF) with localized 2-photon laser illumination through the pupal case. Activated pupae are mounted for long-term live imaging approximately 3-6 h after induction and imaged by confocal microscopy. Alternatively, pupae are maintained until adulthood for examination of the adult wing (approximately 4 days later). **C**. Left: Snapshots of early pupal wings imaged through the pupal case at 10h APF, just before two photon activation of Shine-Gal4. Wings express either DSRF:mScarlet (top) or E-cad:3xmKate2 (bottom) and contain a UAS>TKVQ235D transgene. The activated regions are shown in red. Arrowhead indicates sensory cell used as spatial reference. Middle panels show adult wings 4 days after activation with closeups of induced ectopic veins on the right. **D**. Timelapse images of an induced ectopic vein. Imaging starts 3h after activation. At 26h APF, two clusters of nuclei showing strong downregulation of DSRF:mScarlet can be seen (white arrowheads). By 35h APF, the two clusters appear connected (yellow arrowhead). At 40h APF, the induced vein is regular and smooth. Scale bars: 0.5 mm (C, full adult wings), 0.2 mm (C, adult wing close-ups) and 50 *μ*m (D).

We next aimed to test experimentally our predictions of ectopic transverse vein behaviour. Refinement of ectopic veins has been shown previously in specific mutant background, e.g. in *net* and *plexus* mutants [46–48]. To avoid the confounding effect of these genes on the signaling network, we used optogenetics to trigger the formation of ectopic proveins *in vivo* in an otherwise wild type animal. We built upon the observation that global activation of the Dpp pathway in the pupal wing with Shine-Gal4 is sufficient to cause ectopic venation [49]. Here, localised activation of UAS-TKV^Q253D^ [50, 51], and hence ectopic Dpp signalling was achieved by 2-photon illumination-through the pupal case at 10h APF, to allow time for refinement (Figure 7B). This protocol was sufficient to cause ectopic veins in the adult wing, at locations corresponding to the stimulated regions (Figure 7C and S7D for additional examples). While induction of ectopic veins was highly efficient between L2 and L3 (>80%), we were unable to generate ectopic veins between L3 and L4, suggesting that this region is refractory to vein formation, most likely because of *knot* expression [52]. Remarkably, ectopic transverse proveins induced between L2 and L3 appeared smooth and were 2 to 3 cells wide, irrespective of the size of the induced region (Figure 7C). However, they rarely spanned across the whole space between L2 and L3, despite optogenetic activation across this region (Figure 7C and S7D). To track the refinement of these ectopic transverse proveins, the dynamics of mScarlet:DSRF, an endogenous fusion construct, was recorded (Figure 7D, Movie S8, Methods). The domain of Dpp signalling activation was recognized at 13h APF as zones of reduced mScarlet-DSRF fluorescence, which could be a result of both DSRF downregulation and fluorescence bleaching (Figure S7C). However, from 21 to 26h APF, a further decay in mScarlet:DSRF intensity could be observed, indicating active cell fate regulation (Figure 7D, white arrowheads). From 35 to 40h APF, the wide ectopic transverse provein domain became considerably smoother and regular, eventually achieving the characteristic 2 to 3 cell width seen for endogenous veins (Figure 7D, 40h APF). Although we have not determined why ectopic transverse proveins rarely span the whole domain of activation, our results support the conclusion that the signalling network controlling provein refinement has the intrinsic ability to form coherent and smooth stripes of characteristic width.

## Discussion

Here we have used wing veins of *Drosophila* to determine how a multi-signal network drives the reproducible patterned specification of cell fates within a dynamic tissue. The approximate domains where wing veins are fated to form are recognised in the third instar as broad jagged-edged bands of DSRF repression. Yet, after subsequent wing morphogenesis, veins have a highly reproducible width of alternating 2 and 3 cell diameters and a smooth outline. To track vein refinement and fate adjustments in real time, we first devised a live reporter of DSRF expression. We find that, initially (between 16 and 22h APF), prospective veins narrow as a result of local tissue deformation. Beyond ~22h APF, the role of tissue shape change becomes negligible and cell fate change becomes the dominant feature. Specifically, DSRF-negative cells progressively gain DSRF expression in a pattern that smoothens and thins out the initially crude provein domain. We have analysed how these changes in cell fate are choreographed to ensure precise and robust vein formation. Three mutually dependent signalling pathways have been previously implicated in vein fate specification, those mediated by Dpp, EGFR, and Notch. We confirmed with live reporters that these pathways are active in and around proveins and we resolved their temporal evolution. We found that, during this period of development, Notch acts over several cell diameters while Dpp and Spitz are juxtacrine, as described previously [39, 53].

Notwithstanding this unexpected feature, our experimental results suggest that final vein width is achieved by a balancing out of vein promoting and vein inhibiting influences. To explore this intuition, we developed a mathematical model that incorporates the experimentally measured dynamics of reporter gene expression.

In the model, the actions of Spitz and Dpp are combined in a single vein-promoting signal, while Delta suppresses vein fate via activation of the Notch receptor. Since Spitz, Dpp, and Delta are produced by provein cells, proveins are a source of opposing signals. As the model shows, for vein refinement to take place, it is essential that these signals act at different range (long-range inhibition and short-range activation), as observed experimentally. We find that three key parameters determine the outcome of simulations: *J^A^*, the strength of short range activation of the cell fate, *J^I^*, the strength of vein fate inhibition by Notch, and r, the relative stability of the vein and intervein fate state (how easy it is for a cell to go from the vein to intervein state, compared to moving from the intervein to vein state). We determined the phase space that allows provein domains to reach a reproducible width (Supplementary Theory). We find that cis-inhibition of Notch signalling significantly enlarges the size of the parameter space in which stable veins can occur, providing with a potential function for this feature of Delta/Notch signalling. The model also uncovers a smoothening effect arising from nearest-neighbour activation that formally resembles line tension despite the absence of mechanical effects. Interestingly, increase of mechanical line tension at the vein/intervein boundary relative to vein/vein and intervein/intervein boundaries does occur, but only at late stages, once vein refinement has completed (Figure S3C). Exploring the parameter space, we found that the model predicts that, as nearest-neighbour activation is decreased and at sufficiently low cis-inhibition, the initial stripe prepattern disappears and discrete clusters emerge (Figure S5F). In this limit, our model indeed resembles that proposed to account for the specification of sensory bristles on the fly thorax, where cells exhibit bistability, inhibition over a distance of a few cells, but no nearest neighbour activation [41]. Therefore, a repertoire of patterns can emerge from the relatively simple ingredients of bistability and local activation and inhibition.

As a test of our conclusion that cell-fate adjustments are sufficient to drive vein refinement, we used optogenetics to activate proveins in bands of cells orthogonal to normal longitudinal veins and tracked their evolution in pupal wings. These artificial transverse proveins did refine and often formed normal looking vein structures in adults, at a location and orientation corresponding to the domain of activation. This confirms that the refinement process does not rely on long range tissue rearrangements, i.e., is locally autonomous, most likely driven by the signalling network. For transverse veins, our model recapitulated this behaviour with experimentally derived parameters, albeit imperfectly since they disappeared when simulations were extended for long periods of time. It appears therefore that the refinement network alone may not completely account for the refinement of veins that are orthogonal to the axis of tissue elongation. Moreover, experimental ectopic transverse proveins rarely maintained a connection with both endogenous longitudinal veins even though the artificial transverse proveins were activated across the whole intervein domain. We note in this respect that natural cross veins, while still being Dpp and Notch dependent, form with a distinct mechanism perhaps as an assurance to counter the effect of tissue elongation [15]. Interestingly, myosin contractility has also been implicated in the refinement of the posterior cross vein but not in longitudinal veins, indicating that distinct refinement mechanisms may operate in different veins [54]. These considerations suggest that features not incorporated in our model, such as perhaps planar cell polarity [55] or non-uniform Notch expression [22, 23], contribute to robust vein formation. Nevertheless, the model largely captures the behaviour of natural longitudinal veins and could be seen as a template to understand other progressive cell fate decisions that rely on the interaction between signalling and expression of a cell fate determinant, such as occurs for example in early mammalian embryos [56].

## Acknowledgements

We thank Romain Levayer and Florence Levillayer for kindly sharing the unpublished tub>minicic:neonGreen fly stock with us. S.H and J.-P.V. thank the Crick fly facility for support and the Crick advanced light microscopy facility for help with imaging. We kindly thank Maceo Joseph, Brian Simeth, and Louis Combépine for help with the manual correction of movie segmentation and tracking. We thank Yohanns Bellaïche for insightful comments on the manuscript. S.H. was supported by a long-term EMBO fellowship (ALTF 1156-2015) and a fellowship from the Ciência sem Fronteiras program (CNPq, Brazil). J.-P. V. and G.S. were supported by the Francis Crick Institute which receives its core funding from Cancer Research UK (FC001204, FC001317), the UK Medical Research Council (FC001204, FC001317), and the Wellcome Trust (FC001204, FC001317). M. de G. was supported by a PhD studentship from the Francis Crick Institute. S.C. was supported by a SNSF project grant 200021_197068 to G.S.

## Author contributions

Conceptualisation: S.H., M.d.G., S.C., H.A., J.-P. V., G.S. Experiments: S.H., with support from C.A. and A.H. Theory and simulations: M.d.G., S.C., G.S. Software: M.S. Supervision: J.-P. V., G.S. Writing: S.H., J.-P. V., G.S.

## References

1. Kicheva, A. and J. Briscoe, Developmental Pattern Formation in Phases. Trends Cell Biol, 2015. 25(10): p. 579–591.

2. Green, J.B. and J. Sharpe, Positional information and reaction-diffusion: two big ideas in developmental biology combine. Development, 2015. 142(7): p. 1203–11.

3. Hiscock, T.W., P. Tschopp, and C.J. Tabin, On the Formation of Digits and Joints during Limb Development. Dev Cell, 2017. 41(5): p. 459–465.

4. Amourda, C. and T.E. Saunders, Gene expression boundary scaling and organ size regulation in the Drosophila embryo. Dev Growth Differ, 2017. 59(1): p. 21–32.

5. Chan, C.J., C.P. Heisenberg, and T. Hiiragi, Coordination of Morphogenesis and Cell-Fate Specification in Development. Curr Biol, 2017. 27(18): p. R1024–R1035.

6. Schweisguth, F. and F. Corson, Self-Organization in Pattern Formation. Dev Cell, 2019. 49(5): p. 659–677.

7. Sharrock, T.E. and B. Sanson, Cell sorting and morphogenesis in early Drosophila embryos. Semin Cell Dev Biol, 2020. 107: p. 147–160.

8. Montagne, J., et al., The Drosophila Serum Response Factor gene is required for the formation of intervein tissue of the wing and is allelic to blistered. Development, 1996. 122(9): p. 2589–97.

9. Roch, F., et al., Genetic interactions and cell behaviour in blistered mutants during proliferation and differentiation of the Drosophila wing. Development, 1998. 125(10): p. 1823–32.

10. O’Keefe, D.D., et al., Egfr/Ras signaling regulates DE-cadherin/Shotgun localization to control vein morphogenesis in the Drosophila wing. Dev Biol, 2007. 311(1): p. 25–39.

11. O’Keefe, D.D., et al., Discontinuities in Rap1 activity determine epithelial cell morphology within the developing wing of Drosophila. Dev Biol, 2012. 369(2): p. 223–34.

12. Abouchar, L., et al., Fly wing vein patterns have spatial reproducibility of a single cell. J R Soc Interface, 2014. 11(97): p. 20140443.

13. Blair, S.S., Wing vein patterning in Drosophila and the analysis of intercellular signaling. Annu Rev Cell Dev Biol, 2007. 23: p. 293–319.

14. De Celis, J.F., Pattern formation in the Drosophila wing: The development of the veins. Bioessays, 2003. 25(5): p. 443–51.

15. Antson, H., T. Tonissoo, and O. Shimmi, The developing wing crossvein of Drosophila melanogaster: a fascinating model for signaling and morphogenesis. Fly (Austin), 2022. 16(1): p. 118–127.

16. Guichard, A., et al., rhomboid and Star interact synergistically to promote EGFR/MAPK signaling during Drosophila wing vein development. Development, 1999. 126(12): p. 2663–76.

17. Martin-Blanco, E., et al., A temporal switch in DER signaling controls the specification and differentiation of veins and interveins in the Drosophila wing. Development, 1999. 126(24): p. 5739–47.

18. Sotillos, S. and J.F. De Celis, Interactions between the Notch, EGFR, and decapentaplegic signaling pathways regulate vein differentiation during Drosophila pupal wing development. Dev Dyn, 2005. 232(3): p. 738–52.

19. de Celis, J.F., Expression and function of decapentaplegic and thick veins during the differentiation of the veins in the Drosophila wing. Development, 1997. 124(5): p. 1007–18.

20. Sturtevant, M.A., M. Roark, and E. Bier, The Drosophila rhomboid gene mediates the localized formation of wing veins and interacts genetically with components of the EGF-R signaling pathway. Genes Dev, 1993. 7(6): p. 961–73.

21. Sturtevant, M.A. and E. Bier, Analysis of the genetic hierarchy guiding wing vein development in Drosophila. Development, 1995. 121(3): p. 785–801.

22. de Celis, J.F., S. Bray, and A. Garcia-Bellido, Notch signalling regulates veinlet expression and establishes boundaries between veins and interveins in the Drosophila wing. Development, 1997. 124(10): p. 1919–28.

23. Huppert, S.S., T.L. Jacobsen, and M.A. Muskavitch, Feedback regulation is central to Delta-Notch signalling required for Drosophila wing vein morphogenesis. Development, 1997. 124(17): p. 3283–91.

24. Crozatier, M., et al., Vein-positioning in the Drosophila wing in response to Hh; new roles of Notch signaling. Mech Dev, 2003. 120(5): p. 529–35.

25. Etournay, R., et al., TissueMiner: A multiscale analysis toolkit to quantify how cellular processes create tissue dynamics. Elife, 2016. 5.

26. Gonzalez-Gaitan, M., M.P. Capdevila, and A. Garcia-Bellido, Cell proliferation patterns in the wing imaginal disc of Drosophila. Mech Dev, 1994. 46(3): p. 183–200.

27. Etournay, R., et al., Interplay of cell dynamics and epithelial tension during morphogenesis of the Drosophila pupal wing. Elife, 2015. 4: p. e07090.

28. Aigouy, B., et al., Cell flow reorients the axis of planar polarity in the wing epithelium of Drosophila. Cell, 2010. 142(5): p. 773–86.

29. Ray, R.P., et al., Patterned Anchorage to the Apical Extracellular Matrix Defines Tissue Shape in the Developing Appendages of Drosophila. Dev Cell, 2015. 34(3): p. 310–22.

30. Diaz-de-la-Loza, M.D., et al., Apical and Basal Matrix Remodeling Control Epithelial Morphogenesis. Dev Cell, 2018. 46(1): p. 23–39 e5.

31. Landsberg, K.P., et al., Increased cell bond tension governs cell sorting at the Drosophila anteroposterior compartment boundary. Curr Biol, 2009. 19(22): p. 1950–5.

32. Yu, K., et al., The Drosophila decapentaplegic and short gastrulation genes function antagonistically during adult wing vein development. Development, 1996. 122(12): p. 4033–44.

33. Lee, J.R., et al., Regulated intracellular ligand transport and proteolysis control EGF signal activation in Drosophila. Cell, 2001. 107(2): p. 161–71.

34. Housden, B.E., K. Millen, and S.J. Bray, Drosophila Reporter Vectors Compatible with PhiC31 Integrase Transgenesis Techniques and Their Use to Generate New Notch Reporter Fly Lines. G3 (Bethesda), 2012. 2(1): p. 79–82.

35. Cohen, M., et al., Dynamic filopodia transmit intermittent Delta-Notch signaling to drive pattern refinement during lateral inhibition. Dev Cell, 2010. 19(1): p. 78–89.

36. Hadjivasiliou, Z., G.L. Hunter, and B. Baum, A new mechanism for spatial pattern formation via lateral and protrusion-mediated lateral signalling. J R Soc Interface, 2016. 13(124).

37. Valon, L., et al., Robustness of epithelial sealing is an emerging property of local ERK feedback driven by cell elimination. Dev Cell, 2021. 56(12): p. 1700–1711 e8.

38. Dunipace, L., et al., Autoregulatory feedback controls sequential action of cis-regulatory modules at the brinker locus. Dev Cell, 2013. 26(5): p. 536–43.

39. Gui, J., et al., Coupling between dynamic 3D tissue architecture and BMP morphogen signaling during Drosophila wing morphogenesis. Proc Natl Acad Sci U S A, 2019. 116(10): p. 4352–4361.

40. Yan, S.J., et al., Bistability coordinates activation of the EGFR and DPP pathways in Drosophila vein differentiation. Mol Syst Biol, 2009. 5: p. 278.

41. Corson, F., et al., Self-organized Notch dynamics generate stereotyped sensory organ patterns in Drosophila. Science, 2017. 356(6337).

42. Sprinzak, D., et al., Cis-interactions between Notch and Delta generate mutually exclusive signalling states. Nature, 2010. 465(7294): p. 86–90.

43. Goldstein, R.E., D.J. Muraki, and D.M. Petrich, Interface proliferation and the growth of labyrinths in a reaction-diffusion system. Phys Rev E Stat Phys Plasmas Fluids Relat Interdiscip Topics, 1996. 53(4): p. 3933–3957.

44. He, L., et al., In vivo study of gene expression with an enhanced dual-color fluorescent transcriptional timer. Elife, 2019. 8.

45. Zacharioudaki, E. and S.J. Bray, Tools and methods for studying Notch signaling in Drosophila melanogaster. Methods, 2014. 68(1): p. 173–82.

46. Brentrup, D., et al., Regulation of Drosophila wing vein patterning: net encodes a bHLH protein repressing rhomboid and is repressed by rhomboid-dependent Egfr signalling. Development, 2000. 127(21): p. 4729–41.

47. Baonza, A. and A. Garcia-Bellido, Dual role of extramacrochaetae in cell proliferation and cell differentiation during wing morphogenesis in Drosophila. Mech Dev, 1999. 80(2): p. 133–46.

48. Matakatsu, H., et al., Repression of the wing vein development in Drosophila by the nuclear matrix protein plexus. Development, 1999. 126(23): p. 5207–16.

49. di Pietro, F., et al., Rapid and robust optogenetic control of gene expression in Drosophila. Dev Cell, 2021. 56(24): p. 3393–3404 e7.

50. Nellen, D., et al., Direct and long-range action of a DPP morphogen gradient. Cell, 1996. 85(3): p. 357–68.

51. Wieser, R., J.L. Wrana, and J. Massague, GS domain mutations that constitutively activate T beta R-I, the downstream signaling component in the TGF-beta receptor complex. EMBO J, 1995. 14(10): p. 2199–208.

52. Mohler, J., et al., Activation of knot (kn) specifies the 3-4 intervein region in the Drosophila wing. Development, 2000. 127(1): p. 55–63.

53. Sturtevant, M.A., J.W. O’Neill, and E. Bier, Down-regulation of Drosophila Egf-r mRNA levels following hyperactivated receptor signaling. Development, 1994. 120(9): p. 2593–600.

54. Toddie-Moore, D.J., et al., Mechano-chemical feedback mediated competition for BMP signalling leads to pattern formation. Dev Biol, 2022. 481: p. 43–51.

55. Piscitello-Gómez, R., et al., Core PCP mutations affect short time mechanical properties but not tissue morphogenesis in the <em>Drosophila</em> pupal wing. bioRxiv, 2022: p. 2022.12.09.519799.

56. Frum, T. and A. Ralston, Cell signaling and transcription factors regulating cell fate during formation of the mouse blastocyst. Trends Genet, 2015. 31(7): p. 402–10.

