## Supplementary material for "Signalling-dependent refinement of cell fate choice during tissue remodelling": Methods

### Key Resources Table

| Reagent or resource | Source | Identifier |
|---|---|---|
| Experimental models: Organisms/strains | | |
| *D. melanogaster*: Ecad:3xmKate2 | [1] | NA |
| *D. melanogaster*: DSRF>GFP | This study | NA |
| *D. melanogaster*: $dpy^{ov1}$ | Bloomington Drosophila Stock Center | BDSC #276 |
| *D. melanogaster*: NRE>mcherry | [2] | NA |
| *D. melanogaster*: tub>minicic:neonGreen | Gift from Romain Levayer, based on [3] | NA |
| *D. melanogaster*: brinker>GFP | *brkNFgfp* [4] | NA |
| *D. melanogaster*: $N^{55e11}$ | Bloomington Drosophila Stock Center | BDSC #28813 |
| *D. melanogaster*: ubi>$Gal4^{MagHigh}$ | [5] | NA |
| *D. melanogaster*: UAS>$TKV^{Q253D}$ | Bloomington Drosophila Stock Center | BDSC #36536 |
| *D. melanogaster*: DSRF:mScarlet | This study | NA |
| *D. melanogaster*: DSRF-BAC | This study | NA |
| *D. melanogaster*: Ecad:$GFP^{ki}$ | [6] | NA |
| *D. melanogaster*: Ecad:3xTagRFP | [1] | NA |
| *D. melanogaster*: rho>$DBD^{MagHigh}$ | This study | NA |
| *D. melanogaster*: ubi>$AD^{MagHigh}$ | [5] | NA |
| *D. melanogaster*: UAS-LifeAct:GFP | Bloomington Drosophila Stock Center | BDSC #35544 |
| *D. melanogaster*: act5C>FRT-y+-FRT-Gal4 | Bloomington Drosophila Stock Center | BDSC #3953 |
| *D. melanogaster*: UAS>CD8:mcherry | Bloomington Drosophila Stock Center | BDSC #27391 |
| Antibodies | | |
| anti-DSRF | Active Motif | 39093 |
| anti-pMLC (S19) | Cell Signalling Technology | 3671S |
| anti-Delta | Developmental Studies Hybridoma Bank | C594.9B |
| Recombinant DNA (plasmids) | | |
| DSRF-BAC | This study | Bs42 BAC |
| DSRF>GFP | This study | bs-IOeGFP@NTV-PAX-Cherry |
| DSRF:mScarlet | This study | bs-IOmScarlet@NTV-PAX-Cherry |
| rho>$DBD^{MagHigh}$ | This study | RIV11-rho DBDGal4-HighMag |
| Software and algorithms | | |
| Python programming language | Python Software Foundation | http://www.python.org/ |
| Jupyter notebook | | https://jupyter.org/ |

| | | |
|---|---|---|
| R programming language | | https://www.r-project.org/ |
| RStudio | RStudio | https://www.rstudio.com/ |
| Fiji | [7] | http://fiji.sc |
| Tissue Analyzer | [8] | https://grr.gred-clermont.fr/labmirouse/software/WebPA/ |
| Tissue Miner | [9] | https://github.com/mpicbg-scicomp/tissue_miner |
| Matlab2014a | MathWorks | https://uk.mathworks.com/products/matlab |
| Surface projection algorithm | This study | https://github.com/odinsbane/surface_projection |
| ActiveUnetSegmentation | [10] | https://github.com/FrancisCrickInstitute/ActiveUnetSegmentation https://doi.org/10.5281/zenodo.7380171 |
| Codes used in this study | This study | https://github.com/simonecicolini/Veins_patterning |
| Data used for simulations on wing template | This study | https://zenodo.org/record/7625645#.Y-oULi8w3gh |

### Star Methods

### Resource availability

##### Lead contact

Further information and requests for resources and reagents should be directed to and will be fulfilled by the lead contacts, Jean-Paul Vincent: (experiments) and Guillaume Salbreux: (theory).

##### Materials availability

Flies and plasmids are available from the lead contact, Jean-Paul Vincent.

##### Data and code availability

Simulation codes are available on the github repository: https://github.com/simonecicolini/Veins_patterning and use experimental images stored at https://zenodo.org/record/7625645#.Y-oULi8w3gh. Additional analysis code and raw data are available upon request to the lead contacts.

### Experimental model and subject details

##### Fly stocks and genetics

A list of all the *Drosophila melanogaster* fly stocks used in this study are listed in the Key Resources Table. Flies were grown in standard molasses/cornmeal/yeast food at 25°C. Both female and male individuals were used in all experiments, except for $N^{55e11}$ heterozygous flies (shown in Figure 6G) which were all females, since *Notch* is located in the X chromosome. Light-sensitive flies and crosses involving Shine-Gal4 were kept protected from light in cardboard boxes. In those cases, cross handling, pupae selection and mounting for live imaging were performed in a dark room under an amber light source, which consisted in a regular white light table lamp with three layers of Deep Straw colour filter

LP015 (HT-range, Chris James Lighting Filters) placed over the light source, as described in [5].

#### Method details

##### Molecular biology and transgenesis

The plasmids generated in this study are listed in the Key Resources Table. Synthetised DNA fragments and oligonucleotides were obtained from IDT (Integrated DNA Technologies, Inc.).

The DSRF>GFP transcriptional reporter was generated as described in [11]. In order to recapitulate the endogenous expression of the DSRF locus, the EGFP open reading frame was inserted immediately downstream of the endogenous translation initiation codon (ATG) using CRISPR/Cas9-mediated homologous recombination (Figure S1A). Because such an insertion prevents expression of DSRF from this allele, heterozygous animals displayed a mild ectopic venation phenotype, as reported previously in heterozygous *DSRF* mutants [12, 13]. This phenotype could be rescued by a transgenic Bacterial Artificial Chromosome (BAC) carrying one copy of the *DSRF* locus (DSRF-BAC, described below). DSRF-BAC was kept in the background whenever necessary to ensure normal levels of DSRF activity (Figure S1B). In fixed heterozygous DSRF>GFP expressing wings, the pattern of GFP expression recapitulated that of the endogenous DSRF protein labelled with an anti-DSRF antibody (Figure S1C).

The rescue DSRF-BAC transgene was generated by integrating a 42kb bacterial artificial chromosome (BAC CH321-5G18) containing a copy of the *blistered* genomic sequence on an attP docking site on the third chromosome (attP2 located at 68A4). The integration vector (Bs42) was generated using recombineering-mediated gap repair.

For the DSRF:mScarlet fusion construct, a similar integration vector as described for DSRF>GFP was inserted immediately downstream of the endogenous ATG of *DSRF*, to generate a transcriptional reporter. A N-terminal mScarlet:DSRF fusion was subsequently generated directly in the resulting transgenic flies, by excision of the loxP flanked pax>mcherry marker cassette via Cre-mediated recombination.

To obtain the rho>DBD$^{MagHigh}$ transgene, a *rhomboid* knock-out was first generated by CRISPR/Cas9-mediated recombination to delete 2081bp corresponding to the last three coding exons of *rhomboid,* which were replaced by a attP docking site (153bp upstream of the original exon 2). A attB vector containing a copy of the DNA binding domain of Shine-Gal4 (DBD$^{MagHigh}$, described in [5]) was then inserted in the *rhomboid* attP docking site, allowing authentic expression under the control of the endogenous *rhomboid* promoter.

##### Immunostaining

Whole pupae were fixed in 4% methanol-free formaldehyde (Pierce, Thermo Fisher Scientific, 28906) for 1h and rinsed three times for a total of 30min in PBS + 0.1% Triton-X-

100. Wings were then dissected out of the cuticular sac and incubated in a solution of primary antibodies diluted in PBS + 0.1% Triton-X-100 overnight at 4°C. Samples were then washed three times for a total of 30min in PBS + 0.1% Triton-X-100 and incubated at room temperature for 1.5 h with the secondary antibody solution (1/500) and DAPI, diluted in PBS + 0.1% Triton-X-100. After secondary incubation, the samples were rinsed three times for a total of 30 min in PBS + 0.1% Triton-X-100 and mounted in Vectashield Antifade Mounting Medium with DAPI (Vector Laboratories, H-1200-10). A thin layer of regular nail polish was applied to both sides of the mounting slide to prevent the coverslip from pressurizing and damaging the wings. The primary antibodies used in this study were anti-DSRF (1/500, Active Motif #39093, discontinued), anti-phospho-myosin light chain S19 (1/100, Cell Signalling Technology, #3671S) and anti-Delta extracellular domain (1/100, Developmental Studies Hybridoma Bank, # C594.9B).

**Imaging, optogenetics and laser ablation**

*Long-term live imaging*

Pupae were collected and mounted for live imaging as described previously [14], and the dorsal surface of wings were imaged. Developmental timing was counted from either pre-pupal formation (0h APF) or head eversion (12h APF) and carried out at 25°C. Imaging was performed on an inverted Nikon CSU-W1 Spinning disk microscope equipped with a Prime 95B sCMOS camera (Teledyne Photometrics) and a 40x NA 1.15 Apo LWD Lambda WI objective. Laser lines used were 488nm for green fluorescent protein imaging and 561nm for red fluorescent proteins. Wings were imaged in a single field-of-view, over 71 z-planes (with 1µm spacing between planes) and every 5 minutes, for the whole duration of the experiment (20h on average). Incubation temperature was set to 25°C for all experiments.

*Optogenetics*

The flies used in the optogenetics experiments were timed at 10h APF. Mounting for optogenetic activation was performed as described in [14], but pupae were kept intact (no opening of the pupal case) and illumination was performed through the pupal case. Local Shine-Gal4 activation was performed on a LSM780 NLO inverted confocal microscope (Carl Zeiss) equipped with a Chameleon NIR tuneable two-photon laser (Coherent) and using a 40x NA 1.2 C-Apochromat WI objective. Single-photon imaging with a 561nm laser was used to locate the tissue and define the activation region. Incubation temperature was set to 25°C for all experiments. Two-photon laser power was set to 14.7mW and the activation region was scanned continuously over 6-12 z-planes (so that the nuclear plane of all cells in the activation region was illuminated) for a minimum of 5 minutes and a maximum of 10 minutes, depending on the size of the z-stack. Pixel dwell time was set to 1.53 µs. Following activation, pupae were removed from the slide and the imaging oil was cleaned from the surface using a painting brush embedded in ethanol 70%. Pupae were kept shielded from light at 25°C until adulthood or until 13-15h APF for live imaging. Laser power measurement was performed on the sample plane using a compact power and energy meter console (ThorLabs, #PM100D) fitted with a high power thermal sensor (ThorLabs, #S470C).

*Adult wing mounting and imaging*

Adult flies were preserved and their wings dissected in 100% isopropanol. Wings were mounted on Euparal mounting medium (Anglian Lepidopterist Supplies, ALS). Slides were imaged on a Zeiss Axioplan 2 upright light microscope (Carl Zeiss) equipped with an Axiocam camera and a 5x dry objective.

*Laser ablation*

Laser ablation experiments were performed in flies expressing Ecad:GFP and NRE>mcherry , on an inverted LSM780 NLO confocal microscope (Carl Zeiss) equipped with a Chameleon NIR tuneable two-photon laser (Coherent) and a Plan Apochromat 40x/1.3 Oil DIC M27 objective. The two-photon laser was tuned to 720nm and used at 13% power (250 mW). Single cell-cell junctions were ablated in the centre, at the level of the *adherens* junctions (plane with the strongest Ecad:GFP signal) and across all vein and intervein regions. Intervein cells were determined by the expression of NRE>mcherry. Following laser ablation, single-plane confocal images were immediately acquired every 330ms for a total of 100 frames (33 seconds).

**Image processing, segmentation and tracking**

*Surface projection*

Prior to image analysis, three-dimensional image stacks were converted to 8-bit and projected into 2D planes using a custom-made surface projection algorithm. The E-cad:3xmKate2 channel was used as reference. In brief, for each image stack, a gaussian filter was applied to images to removes features smaller than ~10 $\mu$m. For each x, y values in the image stack, the z-position of the brightest pixel was extracted, to create a height map. The height map was subsequently smoothed to reduce outliers, such as bright pixels located above or below the apical epithelial plane. An implementation of this algorithm is available at https://github.com/odinsbane/surface_projection. The resulting height map was then used to project the original 3D image stacks in 2D, by sampling a set of pixels about the height map values along the z axis.

*Image segmentation*

Projected images were segmented using a U-Net convolutional neural network [15], as described in [10]. To tailor the U-Net neural network to pupal wing images, we included training data obtained from one fully segmented E-cad:3xmKate2-expressing pupal wing movie (1024 x 1024 pixels, 200 frames). The training skeleton was obtained and manually corrected using the Tissue Analyzer Fiji plugin [8]. The trained U-Net model used is available at https://doi.org/10.5281/zenodo.7380171. Following U-Net segmentation, the remaining segmentation errors in the resulting skeletons were manually corrected using Tissue Analyzer.

*Single cell tracking*

Manually-corrected skeletons derived from the U-Net segmentation network were used for single-cell tracking. The tracking algorithm used is described in [10] and is implemented in Matlab. The algorithm outputs a series of RGB images where each cell is given a unique colour ID that is preserved throughout the movie. It also outputs images where cell divisions and suspected tracking errors are highlighted (as described in [10]). These were loaded into Tissue Analyzer, which was used for tracking inspection and manual correction where necessary. The corrected tracking images were input into Tissue Miner [9] to extract individual cell positions (centroids) and lineage relationships. Any remaining segmentation and tracking errors identified in Tissue Miner (cell divisions occurring at late frames and segmentation error appearances) were further corrected using Tissue Analyzer.

*Fluorescence intensity quantification*

Single-cell DSRF>GFP and NRE>mcherry mean intensities were quantified from segmented images using a custom-made algorithm. We determined a height map as described in "Surface projection" and for each pixel obtained the maximum projection of intensities in a 3 pixel-wide region along the z axis, around the detected height. The algorithm then extracted the pixels corresponding to each individual cell using the tracking output RGB images as reference, where each cell is labelled with a unique colour. The pixel-wise GFP or mcherry raw intensity values were summed and averaged over the pixels corresponding to each uniquely coloured region. The measured mean intensity values per cell and per frame were subsequently appended to the Tissue Miner database for further analysis.

To evaluate profiles of DSRF>GFP and NRE>mcherry mean intensities in non-segmented images (Figures 4C, 4D, S6C, S6H-H''), we used maximum intensity projections over the full z-stacks.

#### Data analysis and quantification

*Vein/intervein state classification*

DSRF>GFP mean intensity on each individual cell and at every time point was measured as described above. Cells with a DSRF>GFP mean intensity value above a threshold value were classified as intervein and cells with mean intensity values below the threshold were classified as vein. A threshold value was determined for each individual movie, based on the global distribution of DSRF>GFP mean intensities of all cells and all timepoints. The global intensity distribution was assumed to be a mixture of two gaussians, one corresponding to the vein cell population and the other to the intervein cell population (Figure S3A). The R library *mixtools* (https://rdrr.io/cran/mixtools/) was used to determine the mean and standard deviation of the two gaussians. The threshold was defined as the mean + 1.5 times the standard deviation of the first gaussian. Note that since the intensity distribution in each individual movie might vary, the absolute threshold value can differ between movies.

*Quantification of provein width*

Quantification of provein width was performed using masked provein images. For non-segmented images (Figures 1B and 2B), a Gaussian filter (sigma: 3) was applied to the

DSRF>GFP channel and the global pixel intensity distribution (all pixels over all frames) was determined. A vein/intervein state classification threshold for each individual movie was computed using the intensity distribution, as described above. The classification threshold was then used to binarize the images and generate provein masks, where provein pixels were given a value of 1 and intervein pixels a value of 0. For segmented movies, the areas corresponding to the cells classified as provein were used to create the provein masks. Masked images were rotated to orient proveins horizontally and the width of the provein at every x-coordinate position was computed as the sum of all pixel values at the given x-coordinate. The provein width at each timepoint is given as the mean value across all x-coordinate positions.

*Spatio-temporal alignment of proveins*

Wings from different individuals were registered in time by using cell division dynamics. Based on [16], the time of the last cell division was set to 24h APF in wild-type wings and 26h APF in *dumpy* mutant wings. Time-matched masked provein images (generated as described above) were then spatially aligned by applying a rigid body image registration algorithm to find the best overlap between images.

*Inter-individual provein variability quantification*

Spatio-temporally aligned masked provein images were used to quantify provein variability. First, a reference line passing through to the centre of the aligned proveins was determined by fitting a polynomial function of degree 2 to all images combined. Then, at every x-coordinate, and for each individual provein, the shortest distance between the border of the provein and the reference line was measured. At each x-coordinate, the standard deviation between the measured distances across all proveins was computed (the better the overlap between proveins, the lower the standard deviation). Finally, the standard deviation values between the distances at every x-coordinate were averaged so that a single variability measure was determined at each timepoint.

*Quantification of provein cell count*

Provein cell counts were quantified for both the *initial provein's progeny* and *actual provein* (defined in Figure 3B). For the *initial provein's progeny* cell counts, cells classified as vein in the first frame of the movie and their descendants were tracked and counted at each timepoint. These counts are influenced by cell gains through cell division and loss through delamination, but not by fate changes. For *actual provein* cell counting, at each timepoint, cells that were classified as veins were counted. In addition to incorporating cell divisions and delamination (if any), these counts are also influenced by cell fate changes as cells upregulate or downregulate DSRF>GFP expression.

*Quantification of provein roughness*

Provein roughness was quantified using masked provein images, generated as described above. For each individual provein, a reference line passing through the centre of the provein mask was determined by fitting a polynomial function of degree 2 to the mask. At every x-coordinate, the shortest distance from the border of the provein to the reference line was measured. The x-coordinate values were binned in windows of 50 pixels, which corresponded to approximately 3 cell diameters and the standard deviation between the distances in each of the bins was calculated (the smoother the border, the smaller the standard deviation value). This measure reflects the small-scale roughness of the provein domain (3-cell scale). Finally, to quantify the large-scale roughness of the provein, the standard deviation of the small-scale standard deviation across the whole provein was computed and plotted in Figure 3G.

*Quantification of profiles of DRSF and NRE fluorescence intensity profiles*

To obtain profiles of DSRF>GFP or NRE>mcherry fluorescence as a function of a coordinate going perpendicular to the vein (Figures 4C, 4D, 6D, 6F, S6B, S6C, S6D and S6H-H''), we first determined a line going through the center of the vein at different times; either manually (Figures 4C, 4D, S6C and S6H-H'') or using an automatic procedure as follows (Figures 6D, 6F, S6B and S6D): we select at each time point a group of cells with DSRF>GFP intensity above a threshold to identify the location of vein L3. We then fit a polynomial to the resulting set of vein cell centroids obtained from Tissue Miner. The resulting line is used to define the vein center position.

With the vein center line at hand, we defined a coordinate $z$ perpendicular to the fitted line. To obtain profiles in Figures 4C, 4D, 6D, S6C and S6H-H'', we collected, for a set of pixels around the vein L3, the z-coordinate of the pixel, corresponding to its distance to the line defining the vein position, and its intensity. We then averaged and binned this dataset to obtain a profile of signal intensity as a function of $z$. To obtain the profiles in Figures 6D, 6F, S6B and S6D, for each cell in a region around the vein L3, we collected the $z$ coordinate of the cell, corresponding to the shortest distance between its center and the line defining the vein position, and the average signal intensity on the surface of that cell. We then averaged and binned this dataset to obtain a profile of signal intensity as a function of $z$.

*Quantification of junction recoil velocity*

To quantify recoil velocity following single junction ablation, post-ablation time-lapse movies were used to generate kymographs along each of the ablated junctions in Fiji. The Ecad:GFP channel was used to determine the positions of the two junction vertices at each timepoint by generating fluorescence plot profiles and determining the coordinates of the two intensity peaks. The distance between the two peaks was measured at each timepoint using Fiji. The initial recoil velocity was determined as the slope of a linear function fitted to the first 6 datapoints of the vertex distance data (from 0 to 2s after the ablation). Two-sided t-tests were performed in R to determine statistical significance between the initial recoil velocity distributions of different junction types (border, inside and outside) at the different timepoints.

*Quantification of basal protrusion length*

To observe individual basal protrusions, a flip-out strategy was used to label single cells with a UAS>CD8:mcherry membrane marker. In brief, flies containing a act5C>FRT-y+-FRT-Gal4 flip-out cassette were crossed with hsp70>FLP; UAS>CD8:mcherry flies and their descendants heat-shocked on a water bath at 37°C for 15min at 12hAPF. 3D confocal images were acquired at 20hAPF using a Leica TCS SP5 microscope equipped with a 63x HC PL Apochromat NA 1.4 oil immersion objective and using a z-step size of 0.5µm. The length of individual basal protrusions was manually measured from the cell body to the furthest visible tip using Fiji.


1. Pinheiro, D., et al., *Transmission of cytokinesis forces via E-cadherin dilution and actomyosin flows.* Nature, 2017. **545**(7652): p. 103-107.
2. Housden, B.E., K. Millen, and S.J. Bray, *Drosophila Reporter Vectors Compatible with PhiC31 Integrase Transgenesis Techniques and Their Use to Generate New Notch Reporter Fly Lines.* G3 (Bethesda), 2012. **2**(1): p. 79-82.
3. Valon, L., et al., *Robustness of epithelial sealing is an emerging property of local ERK feedback driven by cell elimination.* Dev Cell, 2021. **56**(12): p. 1700-1711 e8.
4. Dunipace, L., et al., *Autoregulatory feedback controls sequential action of cis-regulatory modules at the brinker locus.* Dev Cell, 2013. **26**(5): p. 536-43.
5. di Pietro, F., et al., *Rapid and robust optogenetic control of gene expression in Drosophila.* Dev Cell, 2021. **56**(24): p. 3393-3404 e7.
6. Huang, J., et al., *From the Cover: Directed, efficient, and versatile modifications of the Drosophila genome by genomic engineering.* Proc Natl Acad Sci U S A, 2009. **106**(20): p. 8284-9.
7. Schindelin, J., et al., *Fiji: an open-source platform for biological-image analysis.* Nat Methods, 2012. **9**(7): p. 676-82.
8. Aigouy, B., D. Umetsu, and S. Eaton, *Segmentation and Quantitative Analysis of Epithelial Tissues.* Methods Mol Biol, 2016. **1478**: p. 227-239.
9. Etournay, R., et al., *TissueMiner: A multiscale analysis toolkit to quantify how cellular processes create tissue dynamics.* Elife, 2016. **5**.
10. Davis, J.R., et al., *ECM degradation in the Drosophila abdominal epidermis initiates tissue growth that ceases with rapid cell-cycle exit.* Curr Biol, 2022. **32**(6): p. 1285-1300 e4.
11. Poernbacher, I., et al., *Lessons in genome engineering: opportunities, tools and pitfalls.* . BioRxiv, 2019: p. 710871.
12. Montagne, J., et al., *The Drosophila Serum Response Factor gene is required for the formation of intervein tissue of the wing and is allelic to blistered.* Development, 1996. **122**(9): p. 2589-97.
13. Roch, F., et al., *Genetic interactions and cell behaviour in blistered mutants during proliferation and differentiation of the Drosophila wing.* Development, 1998. **125**(10): p. 1823-32.
14. Classen, A.K., et al., *Imaging Drosophila pupal wing morphogenesis.* Methods Mol Biol, 2008. **420**: p. 265-75.
15. Çiçek, Ö., et al., *3D U-Net: learning dense volumetric segmentation from sparse annotation*, in *International conference on medical image computing and computer-assisted intervention*. 2016, Springer. p. 424-432.

16. Etournay, R., et al., *Interplay of cell dynamics and epithelial tension during morphogenesis of the Drosophila pupal wing.* Elife, 2015. **4**: p. e07090.
